# Extended Synaptotagmin is a presynaptic ER Ca^2+^ sensor that promotes neurotransmission and synaptic growth in *Drosophila*

**DOI:** 10.1101/141333

**Authors:** Koto Kikuma, Daniel Kim, David Sutter, Xiling Li, Dion K. Dickman

## Abstract

The endoplasmic reticulum (ER) is an extensive presynaptic organelle, exerting important influences at synapses by responding to Ca^2+^ and modulating transmission, growth, lipid metabolism, and membrane trafficking. Despite intriguing evidence for these crucial functions, how presynaptic ER influences synaptic physiology remains enigmatic. To gain insight into this question, we have generated and characterized mutations in the single *Extended Synaptotagmin* (*Esyt*) ortholog in *Drosophila*. Esyts are evolutionarily conserved ER proteins with Ca^2+^ sensing domains that have recently been shown to orchestrate membrane tethering and lipid exchange between the ER and plasma membrane. We first demonstrate that Esyt localizes to an extensive ER structure that invades presynaptic terminals at the neuromuscular junction. Next, we show that synaptic growth, structure, function, and plasticity are surprisingly unperturbed at synapses lacking *Esyt* expression. However, presynaptic overexpression of *Esyt* leads to enhanced synaptic growth, neurotransmission, and sustainment of the vesicle pool during intense levels of activity, suggesting that elevated Esyt at the ER promotes constitutive membrane trafficking or lipid exchange with the plasma membrane. Finally, we find that *Esyt* mutants fail to maintain basal neurotransmission and short term plasticity at elevated extracellular Ca^2+^, consistent with Esyt functioning as an ER Ca^2+^ sensor that modulates synaptic activity. Thus, we identify Esyt as a presynaptic ER Ca^2+^ sensor that can promote neurotransmission and synaptic growth, revealing the first *in vivo* neuronal functions of this conserved gene family.

## INTRODUCTION

The endoplasmic reticulum (ER) is a continuous intracellular organelle with critical but enigmatic roles at synapses. The ER in neurons is involved in diverse functions including synthesis, modification, and trafficking of proteins and lipids, as well as local regulation of Ca^2+^ homeostasis (Renvoise and Blackstone, 2010; Kwon et al., 2016). Indeed, the importance of synaptic ER is further underscored by its involvement in human disease, including hereditary spastic paraplegias (Montenegro et al., 2012; Noreau et al., 2014), amyotrophic lateral sclerosis (Yang et al., 2009; Fasana et al., 2010), Parkinson’s disease (Stutzmann and Mattson, 2011), and Alzheimer’s disease (Zhang et al., 2009; Stutzmann and Mattson, 2011). At presynaptic terminals, recent studies have established important roles for the ER in both membrane trafficking and Ca^2+^ signaling. These studies have revealed that constitutive membrane trafficking is guided through presynaptic ER to the plasma membrane, necessary for the delivery of transmembrane and secreted proteins required for synaptic growth and maintenance (Pfenninger, 2009; Ramirez and Couve, 2011). In addition, presynaptic ER tightly regulates local Ca^2+^ dynamics by orchestrating Ca^2+^ release and sequestration at the ER (Bardo et al., 2006; Kwon et al., 2016). However, it is unclear whether and how presynaptic ER modulates synaptic strength during synaptic activity.

Extended Synaptotagmins (Esyts) are a family of Ca^2+^ sensitive proteins that are attractive candidates to function as ER Ca^2+^ sensors that modulates local Ca^2+^ dynamics. Esyts are defined by the presence of a hydrophobic stretch (HS) followed by a synaptotagmin-like mitochondorial lipid binding protein (SMP) domain and multiple Ca^2+^-binding C2 domains and were recently identified as evolutionarily conserved from yeast (Tricalbin1-3) through mammals (Esyt1-3) (Min et al., 2007; Manford et al., 2012; Herdman and Moss, 2016). Esyts are targeted to the ER by a HS and can mediate tethering of ER-plasma membrane (PM) contact sites to facilitate ER-PM lipid transfer (Giordano et al., 2013; Saheki et al., 2016; Yu et al., 2016; Saheki and De Camilli, 2017), while other functions for Esyts have also been proposed (Jean et al., 2010; Jean et al., 2012; Tremblay et al., 2015). Interestingly, this membrane tethering and lipid transfer activity by Esyt is only activated upon elevated intracellular concentrations of Ca^2+^ suggesting that Esyt is a low affinity Ca^2+^ sensor that may only be activated during specific conditions of elevated Ca^2+^, such as store-operated Ca^2+^ entry and/or neurotransmission (Idevall-Hagren et al., 2015). However, a recent study reported no apparent changes in ER morphology or function at synapses in mutant mice lacking all three *Esyt* isoforms (Sclip et al., 2016), raising questions about what functions, if any, Esyts may have at synapses.

The fruit fly *Drosophila melanogaster* is a powerful model system to elucidate the in vivo functions of *Esyt*. In contrast to mammals, there is a single, highly conserved *Esyt* ortholog. Further, the fly neuromuscular junction (NMJ) enables an array of imaging, electrophysiological, and genetic approaches to illuminate the fundamental roles of genes at synapses. We have therefore generated and characterized the first *Esyt* mutations in *Drosophila* to test the role of Esyt in synaptic growth, function, and plasticity. Specifically, we examined synapses lacking and overexpressing *Esyt* at basal states and under synaptic stress. We find that Esyt localizes to an extensive presynaptic ER network. We also find no significant changes in the levels of synaptic phospholipids, nor do we observe major defects in synaptic growth, function, or plasticity, in *Esyt* mutants. However, we find that Esyt is necessary for proper neurotransmission and short-term synaptic plasticity at elevated extracellular Ca^2+^ levels, consistent with Esyt functioning as a low affinity ER Ca^2+^ sensor. Furthermore, presynaptic overexpression of *Esyt* promotes synaptic growth, transmission, and sustainment of a functional vesicle pool during intense synaptic activity, suggesting that an increased supply of synaptic membrane due to *Esyt* overexpression may facilitate synaptic growth and vesicle biogenesis. Thus, Esyt is a presynaptic ER Ca^2+^ sensor required for transmission at elevated Ca^2+^ levels, which can promote synaptic growth when expressed at elevated levels.

## MATERIALS AND METHODS

### Fly stocks

All *Drosophila* stocks were raised at 25°C on standard molasses food. The *w^1118^* strain is used as the wild-type control unless otherwise noted, as this was the genetic background into which all genotypes and transgenic lines were crossed. The *Drosophila* stocks used in this study are following: *OK6-Gal4* (Aberle et al., 2002), *BG57-Gal4* (Budnik et al., 1996), *UAS-GFP-KDEL* (Dong et al., 2013; Nandi et al., 2014), *UAS-GFP-myc-2xFYVE* (Wucherpfennig et al., 2003), UAS-PH-PLCδ1-GFP (Verstreken et al., 2009; Khuong et al., 2010). The *Esyt*^2^ mutant (*Mi{ET1}Esyt2^MB029221^*), the *Esyt* deficiency (*Df(3R)Exel7357*), and all other stocks were obtained from the Bloomington Drosophila Stock Center unless otherwise noted. Standard crossing strategies and chromosomal balancers were used. Female larvae were used unless otherwise specified.

### Molecular biology

*Esyt* cDNA (RE26910) was obtained as an expressed sequence tag from the *Drosophila* Genomics Resources Center. Full-length *Esyt* cDNA was PCR amplified using the following primers: F: 5’ CGGCGGTACCCAAAATGAGCGATAACAGTC 3′ and R: 5’ CTACATATGAGCCACCGCCCTCGTGCCGTATTTCAG 3’. The PCR products were subcloned into the pACU2 vector (Han et al., 2011). An mCherry or 3xFlag tag were inserted in-frame at the C-terminal end of the pACU2-*Esyt* construct. To generate Esyt^ΔHS^, the *Drosophila Esyt* hydrophobic stretch was identified by the SMART domain online tool (http://smart.embl-heidelberg.de/). *Esyt* coding DNA before and after the identified hydrophobic stretch were separately PCR amplified and ligated into the pACU2-*mCherry* vector using the Gibson Assembly Cloning Kit (New England Biolabs Inc., E5510S). Finally, for cloning of pACU2-*Esyt^D−N^*, the conserved aspartates in each C2 domains were identified and mutated into asparagine (D364N, D374N, D421N, D423N, E429Q for C2A; D517N, D564N for C2B; D746N, D752N for C2C) by synthesizing and inserting a stretch of DNA that covered the entire C2 domain region. All constructs were sequenced to verify accuracy, and injected into the VK18 recombination site on the second chromosome by BestGene Inc (Chino Hill, CA).

*Esyt*^1^ mutants were generated using a CRISPR/Cas9 genome editing strategy as described (Gratz et al., 2013b; Gratz et al., 2013a; Yu et al., 2013; Bassett and Liu, 2014; Beumer and Carroll, 2014; Sebo et al., 2014). Briefly, a target Cas-9 cleavage site in *Esyt* was chosen using the CRISPR optimal target finder (http://tools.flycrispr.molbio.wisc.edu/targetFinder/). The earliest target in the first common exon shared by all putative *Esyt* isoforms without any obvious off target sequences in the *Drosophila* genome was chosen (sgRNA target sequence: 5’GACAAATGGAAACTCAATTG<u>TGG3’</u>, PAM underscored). DNA sequences covering this target sequence were synthesized and subcloned into the pU6-BbsI-chiRNA plasmid (Addgene 45946) at the BbsI restriction enzyme digestion site. To generate the sgRNA, pU6-BbsI-chiRNA was PCR amplified using the following primers: F: 5’ GGCGAATTGGGTACCGGG 3’ and R: 5’ CTGCAGGAATTCGATAAAAAAGCACC 3’ and cloned into the pattB vector (Bischof et al., 2007). The construct was injected into the VK18 insertion sequence and balanced. 20 lines were generated and sequenced to screen for putative disruptions in the *Esyt* locus. 17/20 lines produced a deletion or insertion that led to frameshift mutations. The line which produced the earliest stop codon (K32stop) was chosen for further analyses and named the *Esyt^1^* allele.

### Immunochemistry

Wandering third-instar larvae were dissected in ice cold 0 Ca^2+^ modified HL3 saline (Stewart et al., 1994; Dickman et al., 2005) containing (in mM): 70 NaCl, 5 KCl, 10 MgCl_2_, 10 NaHCO_3_, 115 Sucrose, 5 Trehelose, 5 HEPES, pH 7.2, and immunostained as described (Dickman et al., 2006). Briefly, larvae were washed three times with modified HL3 saline, and fixed in either Bouin’s fixative (Sigma, HT10132-1L) or 4% paraformaldehyde in PBS (Sigma, F8775). Larvae were washed with PBS containing 0.1% Triton X-100 (PBST) and incubated in primary antibodies at 4°C overnight. The larvae were then washed in PBST and incubated in secondary antibodies at room temperature for two hours. Samples were transferred in VectaShield (Vector Laboratories) and mounted on glass cover slides. The following antibodies were used: mouse anti-Bruchpilot (BRP; nc82; 1:100; Developmental Studies Hybridoma Bank; DSHB); affinity-purified rabbit anti-GluRIII (1:2000; (Marrus et al., 2004; Chen et al., 2017), mouse anti-Flag (1:500; F1804; Sigma-Aldrich), guinea pig anti-vGlut (1:2000; (Daniels et al., 2004; Chen et al., 2017)), mouse anti-FasII (1:20; 1D4; DSHB), mouse anti-GFP (1:1000; 3e6; Thermo Fisher Scientific). Alexa Fluor 647-conjugated goat anti-HRP (Jackson ImmunoResearch) was used at 1:200. Donkey anti-mouse and anti-rabbit Alexa Fluor 488-, Cy3, and Rhodamine Red X secondary antibodies (Jackson ImmunoResearch) were used at 1:400.

### Confocal imaging and analysis

Larval muscles 4 of abdominal segments A2 and A3 were imaged on a Nikon A1R resonant scanning confocal microscope using a 100× APO 1.4NA oil immersion objective with NIS Elements software as described (Chen et al., 2017). The fluorescence signals were excited by separate channels with laser lines of 488 nm, 561 nm, and 637 nm. Images were acquired using identical settings optimized for signal detection without saturation of the signal for all genotypes within an experiment. The general analysis toolkit in the NIS Elements software was used to quantify bouton number, BRP and GluRIII puncta number, and density by applying intensity thresholds on each of the three channels. For live imaging of Esyt^mChe^, wandering third-instar larvae were dissected, washed, and incubated in Alexa Fluor 647-conjugated goat anti-HRP in 0 Ca^2+^ modified HL3 at 1:200 for 5 min. The samples were then washed and covered in 0 Ca^2+^ modified HL3 saline and mounted on glass cover slides. Images were acquired and analyzed as described above.

FM1-43 experiments were performed as described (Dickman et al., 2005). Briefly, larvae were dissected in ice-cold 0 Ca^2+^ modified HL3 and washed, then stimulated for 10 min with a modified HL3 solution containing 90 mM mM KCl and 10 μM FM1-43 (Molecular Probes, Eugene, Oregon). Larvae were then washed in 0 Ca^2+^ saline before imaging. Images were acquired using a Nikon A1R confocal microscope using a 60x APO 1.0NA water immersion objective and imaged as described above. The general analysis toolkit in the NIS Elements software was used to quantify the mean intensity by applying intensity thresholds.

### Western blotting

Third-instar larval CNS extracts (50 animals of each genotype) and adult heads (7 of each genotype) were homogenized in ice cold lysis buffer (10 mM HEPES + 150 mM NaCl, pH 7.4), mixed with an EDTA-free protease inhibitor cocktail (Roche), and run on 4–12% Bis Tris Plus gels (Invitrogen). After blotting onto PVDF membrane (Novex) and incubation with 5% nonfat milk in TBST (10 mM Tris, pH 8.0, 150 mM NaCl, 0.5% Tween 20) for 60 min, the membrane was washed once with TBST and incubated with anti-Esyt (1:2000) and anti-actin (1:2000; JLA20, DSHB) antibodies overnight at 4C. Membranes were washed and incubated with a 1:5000 dilution of horseradish peroxidase-conjugated secondary antibodies (Jackson ImmunoResearch) for 1 h. Blots were washed with TBST and developed with the ECL Plus Western Blotting system (HyGLO). To generate Esyt polyclonal antibodies, a peptide antigen was synthesized consisting of amino acids 799–816 of Esyt (CTQTGLNSWFDLQPEIRHE). This peptide was conjugated to KLH and injected into two rabbits (Cocalico Inc, Pennsylvania). The rabbit immunosera was affinity purified and used at 1:2000.

### Electron microscopy

EM analysis was performed as described (Atwood et al., 1993). Wandering third-instar larvae were dissected in Ca2+-free HL3, then fixed in 2.5% glutaraldehyde/0.1M cacodylate buffer at 4C. Larvae were then washed in 0.1M cacodylate buffer. The whole mount of body wall musculature were placed in 1% osmium tetroxide/0.1M cacodylate buffer at room temperature for 1hr. After washing, larvae were then dehydrated in Ethanol. Samples were cleared in propylene oxide and infiltrated with 50% Eponate 12 in propylene oxide overnight. The following day, samples were embedded in fresh Eponate 12. Electron micrographs were obtained on a Morgagni 268 transmission electron microscope (FEI, Hillsboro, OR). The junctional region was serially sectioned at a thickness of 60–70 nm. The sections were stained in 2% uranyl acetate for 3 minutes, washed briefly 3× in distilled water, stained in Reynolds lead citrate for 1 minute, washed briefly 3× in dIstilled water and dried. Sections were mounted on Formvar coated single slot grids. Larval muscles 6/7 of abdominal segments were viewed at a 23,000× magnification, and recorded with a Megaview II CCD camera. Images were analyzed blind to genotype using the general analysis toolkit in the NIS Elements software.

### Electrophysiology

All dissections and recordings were performed in modified HL3 saline with 0.4 CaCl_2_ (unless otherwise specified). Larvae were dissected and washed several times with modified HL3 saline. Sharp electrode intracellular recordings (electrode resistance between 10–35 MΩ) were performed on muscles 6 of abdominal segments A2 and A3 as described (Chen et al., 2017). Recordings were acquired using an Axoclamp 900A amplifier, Digidata 1440A acquisition system and pClamp 10.5 software (Molecular Devices). Electrophysiological sweeps were digitized at 10 kHz, and filtered at 1 kHz. Miniature excitatory postsynaptic potentials (mEPSPs) were recorded for one min in the absence of any stimulation. Excitatory postsynaptic potentials (EPSPs) were recorded while cut motor axons were stimulated using an ISO-Flex stimulus isolator (A.M.P.I.). Intensity was adjusted for each cell, set to consistently elicit responses from both neurons innervating the muscle segment, but avoiding overstimulation. Data were analyzed using Clampfit (Molecular devices), MiniAnalysis (Synaptosoft), Excel (Microsoft), and SigmaPlot (Systat) software. Average mEPSP, EPSP, and quantal content were calculated for each genotype with and without corrections for nonlinear summation (Martin, 1955). Recordings were rejected when muscle input resistance (R_in_) was less than 3 MΩ, and resting membrane potential (V_rest_) was above −60 mV or if either measurement deviated by more than 10% during the course of the experiment. To acutely block postsynaptic receptors, larvae with intact motor nerves were dissected and incubated with or without philanthotoxin-433 (PhTx; Sigma; 20 μM) for 10 min. Larvae were resuspended in modified HL3, and motor nerves were cut as described (Frank et al., 2006; Dickman and Davis, 2009). For TEVC, muscles were clamped to −70 mV and stimulus train of 4 EPSC pulses were evoked with a 60 Hz, 0.5 msec stimulus duration, while recording in 3 mM Ca^2+^ modified HL3. Short-term plasticity was estimated by dividing the fourth EPSC with the first EPSC. Data were analyzed as described above.

### Statistical Analysis

All data are presented as mean +/−SEM. Data were compared using either a one-way ANOVA and tested for significance using a 2-tailed Bonferroni post-hoc test, or using a Student’s t-test (where specified) with Graphpad Prism or Microsoft Excel software, and with varying levels of significance assessed as p<0.05 (*), p<0.01 (**), p<0.001 (***), ns=not significant. Quantal content was calculated for each individual recording using the equation QC = EPSP/mEPSP. The QC was corrected for non-linear summation for the Ca^2+^-cooperativity analysis, using the equation QC_corrected_ = (EPSP/mEPSP)(1-EPSP/V0)^−1^ where V0 = (reversal potential – resting potential) (Martin, 1955). Full statistical details and information can be found in Table 1.

### Data availability

Fly stocks are available upon request. Table 1 contains genotypes used and full statistical details for each figure.

## RESULTS

### Generation of null mutations in *Drosophila Esyt*

The rodent genome encodes three *extended synaptotagmin* isoforms (*Esyt1, Esyt2*, and *Esyt3*; (Min et al., 2007)). *Esyt1* encodes five C_2_ domains, while *Esyt2* and *Esyt3* each contain three (Fig 1B). In contrast, the *Drosophila* genome encodes a single *Esyt* ortholog, which is most similar to mouse *Esyt2* by amino acid sequence (54% similar, 34% identical). There are four predicted *Esyt* isoforms based on expressed sequence tags (www.flybase.org). However, only one transcript (*Esyt2-RB*) appears to be the major isoform based on the expression profile. RNA-seq data suggests *Esyt* is expressed in embryonic stages after 10h through adults in all tissues examined, consistent with *Esyt* being ubiquitously expressed. We generated null mutations in the *Drosophila Esyt* gene using CRISPR/Cas-9 genome editing technology (Gratz et al., 2013b; Gratz et al., 2013a; Yu et al., 2013; Bassett and Liu, 2014; Beumer and Carroll, 2014; Sebo et al., 2014). Screening of 20 lines with active CRISPR mutagenesis led to 17 with independent deletions or insertions with predicted frameshift mutations in the *Esyt* open reading frame (see methods). We chose one such mutation for further study, *Esyt^1^*, which is predicted to generate a stop codon at position 32, truncating the Esyt protein before the hydrophobic stretch (Fig 1A and 1B). In addition, we obtained a separate *Esyt* mutation (*Esyt*^2^), containing a MiMIC transposon insertion in the first coding intron. This transposon has a gene trap cassette (Venken et al., 2011; Nagarkar-Jaiswal et al., 2015) that is predicted to truncate the *Esyt* transcript by introducing a stop codon at the second amino acid (Fig 1A and 1B). We generated a polyclonal antibody against a C-terminal stretch of the *Drosophila* Esyt protein, which recognized an immunoblot band at ~90 kDa (Fig 1C and 1D), consistent with the predicted molecular mass of Esyt. We confirmed that Esyt is expressed in the adult brain and larval CNS, and that *Esyt^1^* and *Esyt^2^* are protein null mutations by immunoblot analysis (Fig 1C and 1D). Finally, we generated a series of transgenic *Esyt* constructs under *UAS* control for further analysis (Fig 1B). We engineered a full length *Esyt* transgene (*Esyt2-RB* isoform) tagged with both mCherry and 3×Flag tags (Esyt^mCh^), and a separate transgene tagged with only a 3xFlag tag (Esyt^Flag^). For structure/function studies, we also generated an *Esyt* transgene without the conserved hydrophobic stretch (Esyt^ΔHS^), predicted to disrupt membrane targeting of Esyt (Giordano et al., 2013), and specifically mutated each C2 domain to render them unable to bind to Ca^2+^ (Esyt^D−N^; see methods). Using these reagents, we went on to determine the presynaptic localization of Esyt.

**Figure 1:**
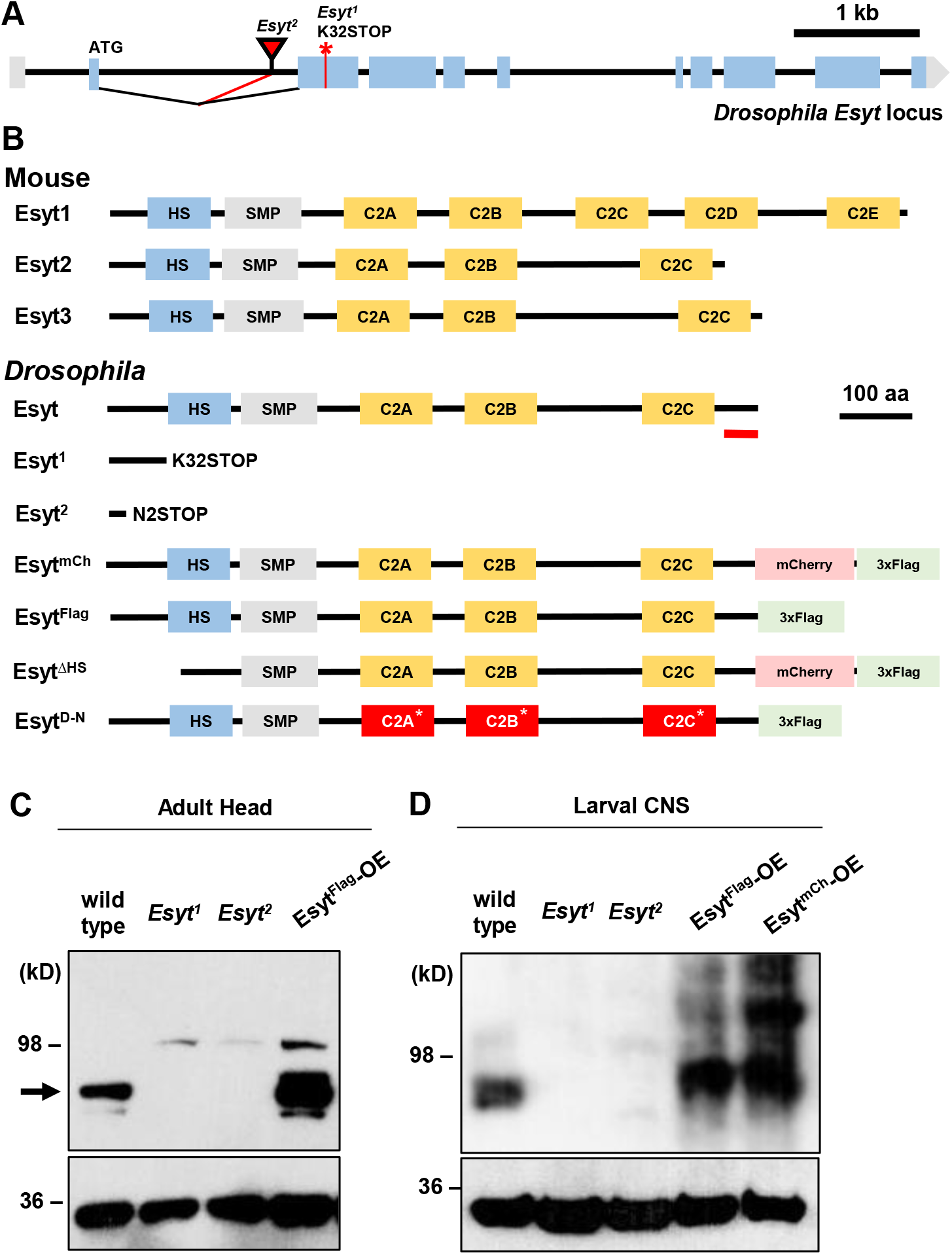
Generation and analysis of null mutations in *Drosophila Esyt*. (**A**) Schematic of the *Drosophila Esyt* locus. The CRISPR induced early stop codon in *Esyt^1^* (red asterisk) and the MiMIC transposon insertion site of *Esyt^2^* (red triangle) are shown. (**B**) Schematic of the mouse Esyt protein structure aligned with the *Drosophila* homolog. The Esyt proteins contain a hydrophobic stretch (HS), synaptotagmin-like-mitochondrial lipid-binding protein (SMP), followed by C2 domains (C2). The Esyt domain structure is highly conserved between *Drosophila* to mice. *Esyt^1^* and *Esyt^2^* mutations truncate the open reading frame before the HS domain. Red line indicates the antigen the antibody was raised against. The structure of *Esyt* transgenes are also shown. Western blot analyses of Esyt in adult heads (C) and larval CNS (D) demonstrates the nervous system expression of Esyt and confirms both *Esyt^1^* and *Esyt^2^* are protein nulls. Neuronal overexpression of *Esyt* (Esyt^Flag^-OE: *w;OK6-Gal4/UAS-Esyt^Flag^* and Esyt^mCh^-OE: *w;OK6-Gal4/UAS-Esyt^mCh^*) results in elevated levels and increased molecular mass observed for the tagged transgene, as expected. Anti-Actin was used as loading control.

### The hydrophobic stretch anchors Esyt to axonal ER

In neurons, the ER is an extensive network present in the somatic perinucleus as well as in distal dendrites and axons (Terasaki et al., 1994; Spacek and Harris, 1997; Meldolesi, 2001; Verkhratsky, 2005; Bardo et al., 2006; Wang et al., 2007; Blackstone et al., 2011; O’Sullivan et al., 2012; Wong et al., 2014; Summerville et al., 2016). Given that ER proteins can have uniform or heterogeneous localization in the elaborate ER network (Chang and Liou, 2016), we sought to determine the subcellular localization of *Drosophila* Esyt at presynaptic terminals. We were unable to obtain specific immunolabeling against endogenous Esyt using the antibody we generated. Therefore, we expressed tagged *Esyt* constructs in motor neurons. Esyt^mCh^ trafficked to presynaptic terminals, where it invaded synaptic boutons and co-localized with an established ER marker, the ER retention signal KDEL fused to GFP (*GFP-KDEL*; Fig 2A; (Dong et al., 2013; Nandi et al., 2014)). Next, we tested whether Esyt localization and trafficking to ER was dependent on the hydrophobic stretch (HS) domain and on Ca^2+^ binding to the C2 domains. Previous studies have shown that the HS domain tethers Esyt to the ER, while deletion of the HS domain shifts Esyt localization to the cytosol and plasma membrane (Min et al., 2007; Giordano et al., 2013). We therefore expressed the mCherry-tagged Esyt construct lacking the HS domain in neurons (Esyt^ΔHS^; see Fig 1B and methods). As expected, the Esyt^ΔHS^ signal was no longer restricted to the axonal ER, and instead filled the presynaptic terminal, indicating a shift to cytosolic localization (Fig 2B). We next tested the requirement of Ca^2+^ binding for Esyt trafficking and localization by expressing Esyt^D−N^, which lacks the negatively charged amino acids in each C2 domain required for Ca^2+^ binding. Interestingly, we were unable to detect any Esyt^D−N^ signal at synaptic terminals (Fig 2C and 2D). Instead, most of the Esyt^D−N^ signal was restricted to the cell body (data not shown). This indicates that trafficking of Esyt to synaptic terminals requires the ability to bind Ca^2+^. However, we cannot exclude the possibility that the D-N mutations might have resulted in misfolding of the protein, potentially disrupting trafficking or stability. We also found that expression of the Esyt^D−N^ transgene led to embryonic lethality when driven pan-neuronally or in muscle. This is not unexpected, as others have observed that Synaptotagmin expression with similar mutations to C2 domains acquired a lethal toxicity (Mackler et al., 2002). Together, these data demonstrate that *Drosophila* Esyt is localized to presynaptic ER structures through the HS domain and that Ca^2+^ binding to Esyt appears to be required for Esyt trafficking in neurons.

**Figure 2:**
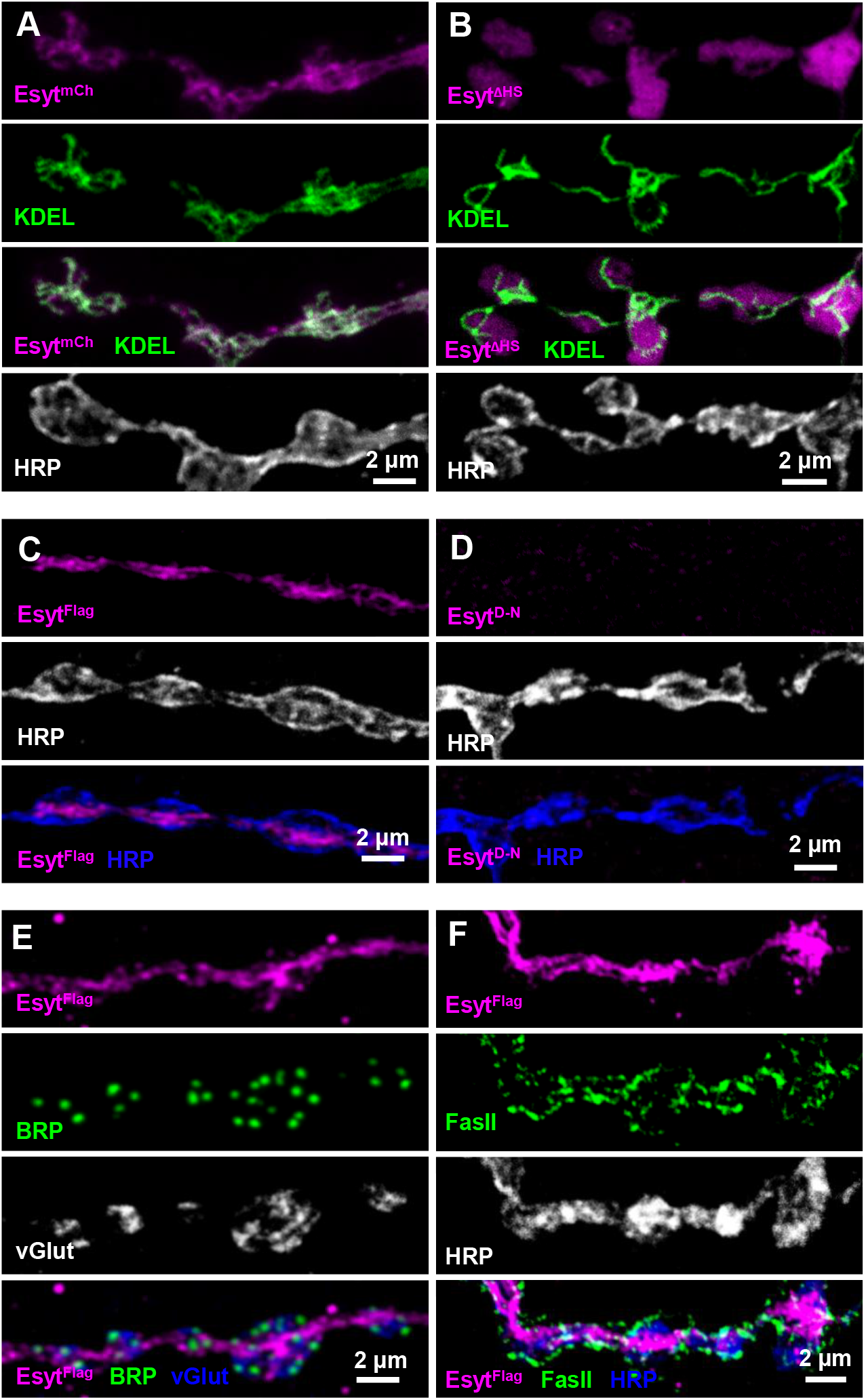
The hydrophobic stretch is necessary to localize Esyt to axonal ER. (**A**) Representative images of third-instar larval NMJ with motor neuron expression of an mCherry-tagged *Esyt* transgene (mCherry; magenta) and a GFP-tagged ER retention signal KDEL (GFP; green; *w;OK6-Gal4/UAS-Esyt^mCh^;UAS-GFP-KDEL*), immunolabeled with the neuronal membrane marker HRP (white/blue). Esyt-mCherry colocalizes with axonal ER labeled by GFP-KDEL. (**B**) Deletion of the hydrophobic stretch from *Esyt-mCherry (w;OK6-Gal4/UAS-Esyt^ΔHS^;UAS-GFP-KDEL*) results in a failure to localize to the ER, instead acquiring a cytosolic distribution. (**C**) Representative NMJ images of a Flag-tagged Esyt transgene (Esyt^Flag^: *w;OK6-Gal4/UAS-Esyt^Flag^*) expressed in motor neurons (anti-Flag; magenta) and labeled with HRP (white/blue). Esyt-Flag traffics to the presynaptic terminals similarly to Esyt-mCherry. (**D**) Mutations in an *Esyt^Flag^* transgene that prevent Ca^2+^ binding to C2 domains (Esyt^D−N^: *w;OK6-Gal4/UAS-Esyt^D−N^*) no longer traffics to presynaptic terminals. (**E**) Axonal ER structures labeled with Esyt-Flag (anti-Flag; magenta) is shown relative to active zones (BRP; green) and synaptic vesicle structures (vGlut; white/blue). (**F**) Axonal ER labeled by Esyt-Flag is shown co-labeled with a peri-active zone marker (FasII; green) and a neuronal membrane marker (HRP; white/blue).

Given that Esyt localizes to axonal ER, we examined the morphology of the ER network at synaptic terminals with loss or overexpression of *Esyt*. Expression of *GFP-KDEL* alone in motor neurons labeled an extensive presynaptic network extending throughout synaptic boutons, as observed by others (data not shown; (Summerville et al., 2016)). This network did not appear to be perturbed in *Esyt* mutants, nor with overexpression (data not shown). These results suggest that ER structure and elaboration is not dependent on *Esyt* expression. Lastly, we found that the ER network labeled by Esyt is localized near, but distinct from, other synaptic compartments including active zones, synaptic vesicle pools, periactive zone regions, and the neuronal plasma membrane (Fig 2E and 2F). Thus, Esyt is localized to the presynaptic ER network and present near areas of synaptic vesicle fusion and recycling at presynaptic terminals where it could, in principle, modulate synaptic structure and function.

### Synaptic phospholipid balance does not require *Esyt*

Given that *Esyt* has been implicated in phospholipid transfer and homeostasis in non-neuronal cells, we next sought to determine whether the level and distribution of phospholipids at presynaptic terminals was altered in *Esyt* mutants and/or with overexpression of *Esyt* in motor neurons (Esyt-OE). The phospholipid phosphatidylinositol 4,5-bisphosphate (PI(4,5)P_2_) plays crucial roles at presynaptic terminals, regulating synaptic protein-protein interactions, ion channel biophysics, neurotransmission, and synaptic vesicle trafficking (De Camilli et al., 1996; Lemmon, 2003; Di Paolo et al., 2004; Di Paolo and De Camilli, 2006; Slabbaert et al., 2012; Lauwers et al., 2016). To determine levels of PI(4,5)P2, we expressed a fluorescently tagged pleckstrin homology domain of phospholipase C-δ1 (PLCδ1-PH-GFP that specifically labels PI(4,5)P2 (Verstreken et al., 2009; Khuong et al., 2010; Chen et al., 2014). Expression of this transgene in motor neurons revealed specific labeling of the plasma membrane at presynaptic terminals, consistent with the expected distribution of PI(4,5)P2 (Fig 3A). When PLCδ1-PH-GFP was expressed in *Esyt* mutants, we were unable to detect any difference in intensity or distribution, nor did PLCδ1-PH-GFP levels change with Esyt-OE (Fig 3A and 3B). Thus, we find no evidence that PI(4,5)P2 levels or distribution is altered at synapses with gain or loss of *Esyt* expression.

**Figure 3:**
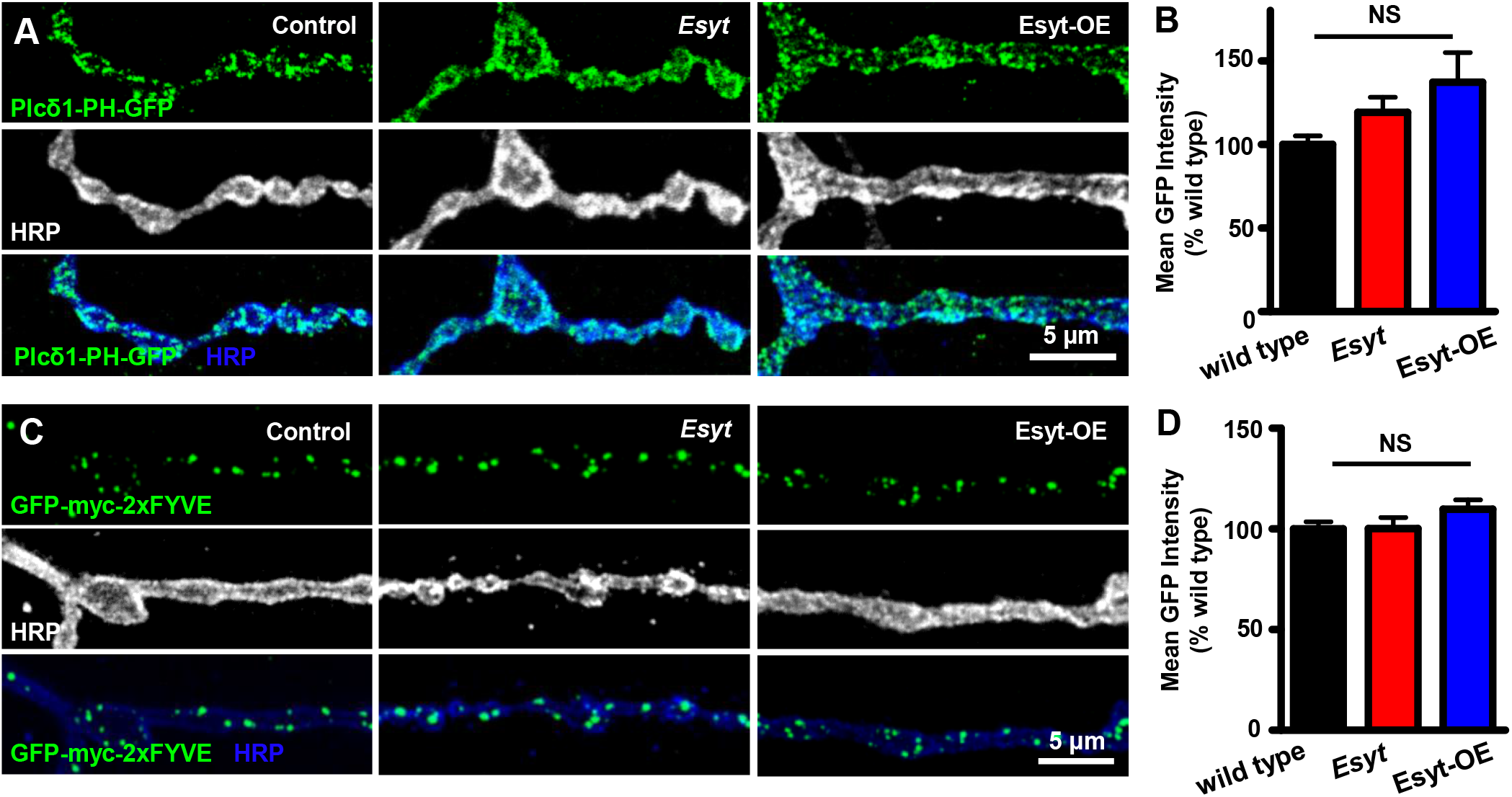
PI(4,5)P_2_ and PI(3)P phospholipid levels are unchanged at presynaptic terminals in *Esyt* mutants and Esyt-OE. (**A**) Representative NMJ images of PI(4,5)P_2_ labeled with PLCδ-PH-GFP (GFP; green) and HRP (white/blue) in control (*w;OK6-Gal4/+;UAS-PLCδ-PH-GFP/+), Esyt* mutants (*w;OK6-Gal4/+;Esyt^1^/Esyt2,UAS-PLCδ-PH-GFP*), and Esyt-OE (w; *OK6-Gal4/UAS-Esyt^mCh^;UAS-PLCδ1-PH-GFP/+*). (**B**) Quantification of mean GFP intensity levels of the indicated genotypes. (**C**) Representative images of PI(3)P distribution at the NMJ labeled by GFP-myc-2xFYVE in control (*w;OK6-Gal4/UAS-GFP-myc-2xFYVE), Esyt* mutants (*w;OK6-Gal4/UAS-GFP-myc-2xFYVE;Esyt^1^/Esyt^2^*), and Esyt-OE (*w;OK6-Gal4/UAS-GFP-myc-2xFYVE;UAS-Esyt^Flag^/+*). (**D**) Quantification of mean GFP intensity levels of the indicated genotypes. Error bars indicate ±SEM. One-way analysis of variance (ANOVA) test was performed, followed by a Tukey’s multiple-comparison test. NS=not significant, p>0.05. Detailed statistical information for represented data (mean values, SEM, n, p) is shown in Table 1.

In addition to the plasma membrane, the ER also associates with early and late endosomal structures (Rowland et al., 2014; Raiborg et al., 2015; Eden, 2016; Phillips and Voeltz, 2016). ER-endosome contact sites modulate endosomal structure and function, contributing to synaptic growth and vesicle trafficking (Wucherpfennig et al., 2003; Rowland et al., 2014; Raiborg et al., 2015). Thus, we considered the possibility that Esyt may regulate lipid transfer or otherwise influence endosomes at synapses. In particular, we focused on early endosomes known to be involved in synaptic vesicle trafficking and recycling. These early endosomes are specifically labeled by the small GTPase Rab5 and are enriched in the phospholipid phosphatidylinositol-3-phosphate (PI(3)P) (Wucherpfennig et al., 2003; Rodal et al., 2011; Slabbaert et al., 2012). The FYVE finger domain of the Rab5 effectors EEA1 and Rabenosyin-5 bind specifically to PI(3)P, which is an established marker for early endosomes (Stenmark et al., 1995; Simonsen et al., 1998; Lawe et al., 2000; Wucherpfennig et al., 2003). To test if PI(3)P levels and/or distribution are dependent on *Esyt* expression, we expressed a GFP-fused FYVE domain transgene (*GPF-myc-2xFYVE*) in *Esyt* mutants and Esyt-OE. GFP-myc-2xFYVE expression in controls labeled a punctate structure in presynaptic boutons, as expected (Fig 3C). However, we observed no differences in the intensity of GFP-myc-2xFYVE in *Esyt* mutants and Esyt-OE compared to control (Fig 3C and 3D). Thus, we find no evidence that Esyt is involved in phospholipid balance, transfer, or distribution at presynaptic terminals.

### Presynaptic overexpression of *Esyt* promotes synaptic growth and transmission

Axonal ER plays a critical role in synaptic growth and neurotransmission in mammals and *Drosophila* (Verkhratsky, 2005; Wang et al., 2007; Pfenninger, 2009; Renvoise and Blackstone, 2010; Blackstone et al., 2011; O’Sullivan et al., 2012; Petkovic et al., 2014; Wong et al., 2014; Kwon et al., 2016; Summerville et al., 2016; de Juan-Sanz et al., 2017). Indeed, recent studies have demonstrated that loss of the genes *atlastin* or *reticulon2*, involved in ER membrane fusion and tube formation, respectively, leads to axonal ER fragmentation and results in increased synaptic growth and decreased neurotransmitter release at the *Drosophila* NMJ (Wang et al., 2007; Blackstone et al., 2011; O’Sullivan et al., 2012; Summerville et al., 2016). We therefore sought to determine to what extent *Esyt* expression contributes to synaptic development and neurotransmission. First, we quantified synaptic growth by immunostaining NMJs with antibodies that recognize neuronal membrane (HRP), active zones (BRP), and postsynaptic glutamate receptors (DGluRIII). *Esyt* mutants exhibited no significant differences in the number of synaptic boutons, nor in the number or density of active zones or glutamate receptor clusters at the NMJ (Fig 4A, 4B, 4D, and 4E). However, overexpression of two independent *Esyt* transgenes in motor neurons, Esyt^mCh^ and Esyt^Flag^, revealed a ~40% increase in synaptic growth, including increased neuronal membrane surface area and total number of active zones per NMJ (Fig 4A-4D). Thus, while loss of *Esyt* has no apparent impact on synaptic growth or structure, elevated levels of *Esyt* in motor neurons promotes synaptic growth at the NMJ.

**Figure 4:**
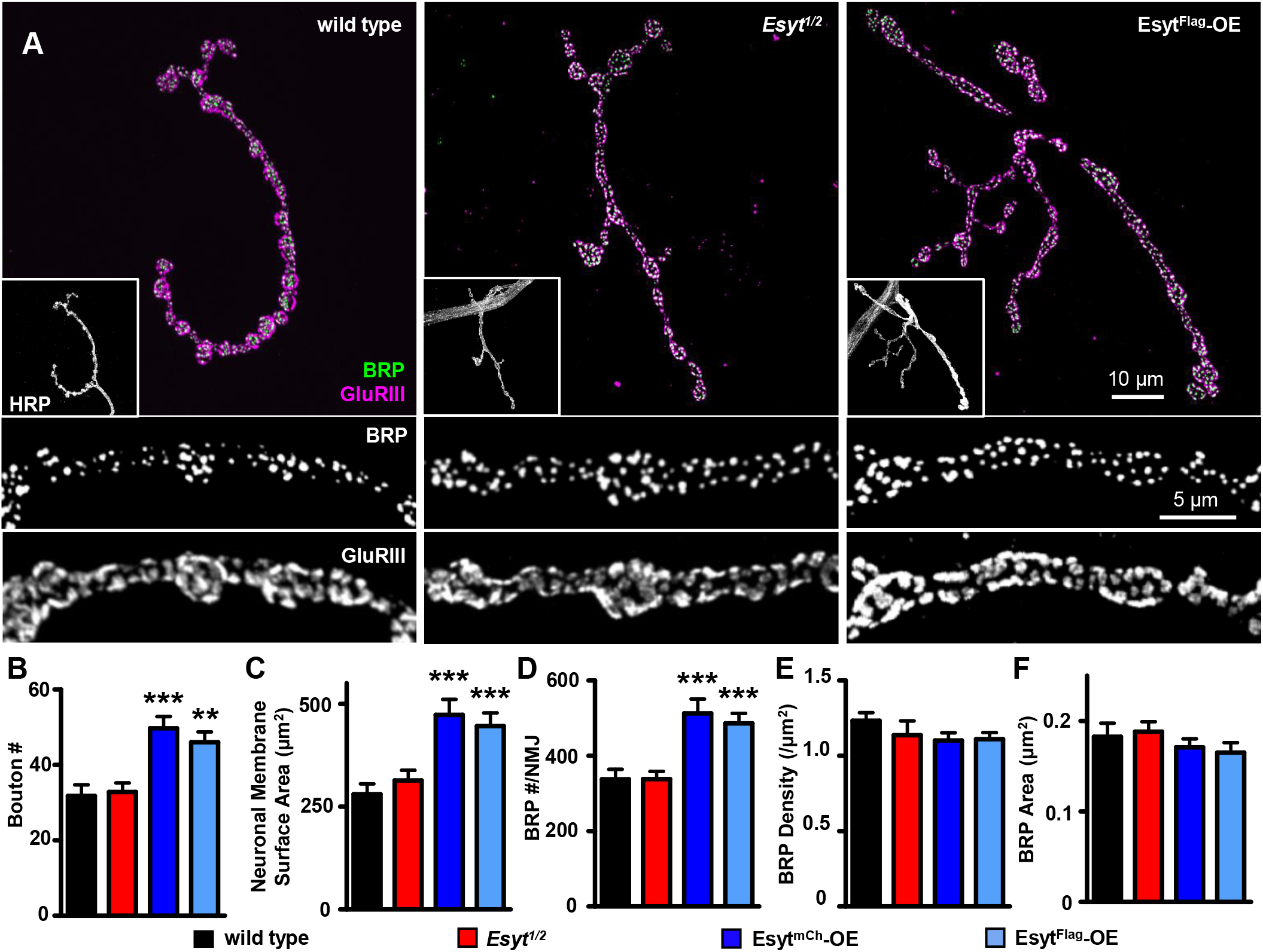
Presynaptic overexpression of *Esyt* promotes synaptic growth. Representative NMJ images of wild type (*w^1118^*), *Esyt* mutants (*Esyt^1/2^: w;Esyt^1^/Esyt^2^*), and Esyt-OE (Esyt^mCh^-OE: *w;OK6-Gal4/UAS-Esyt^mCh^* and Esyt^Flag^-OE: *w;OK6-Gal4/UAS-Esyt^Flag^*) immunostained for anti-BRP (green), anti-GluRIII (magenta), and anti-HRP (white; insert). Bottom panels: BRP and GluRIII images at higher magnification. Quantification of bouton number per NMJ (**B**), neuronal membrane surface area (**C**), total BRP puncta number per NMJ (**D**), BRP density (**E**), and BRP area (**F**) of the indicated genotypes. Note that Esyt-OE results in increased bouton number and a corresponding increase in membrane and BRP number. Error bars indicate ±SEM. One-way ANOVA test was performed, followed by a Tukey’s multiple-comparison test. **p 0.01; ***p 0.001. Detailed statistical information for represented data (mean values, SEM, n, p) is shown in Table 1.

Given that Esyt-OE promotes synaptic growth, we next assessed whether parallel changes in neurotransmission are observed with loss or enhanced expression of *Esyt*. We observed no significant differences in synaptic physiology at lowered extracellular Ca^2+^ (0.4 mM) in *Esyt* mutants compared with controls (Fig 5A-D). In particular, there were no significant changes in mEPSP amplitude, EPSP amplitude, or quantal content (Fig 5A-5D). However, in these conditions, Esyt-OE exhibited a mild increase in EPSP amplitude and quantal content, without any significant difference in mEPSP amplitude (Fig 5A-5D). The increased growth observed in Esyt-OE may contribute to this enhancement of synaptic strength, although clearly there is less change than might be predicted by the increase in growth. Thus, while *Esyt* mutants have no obvious defects in synaptic growth, structure, or transmission, Esyt-OE promotes growth and function.

**Figure 5:**
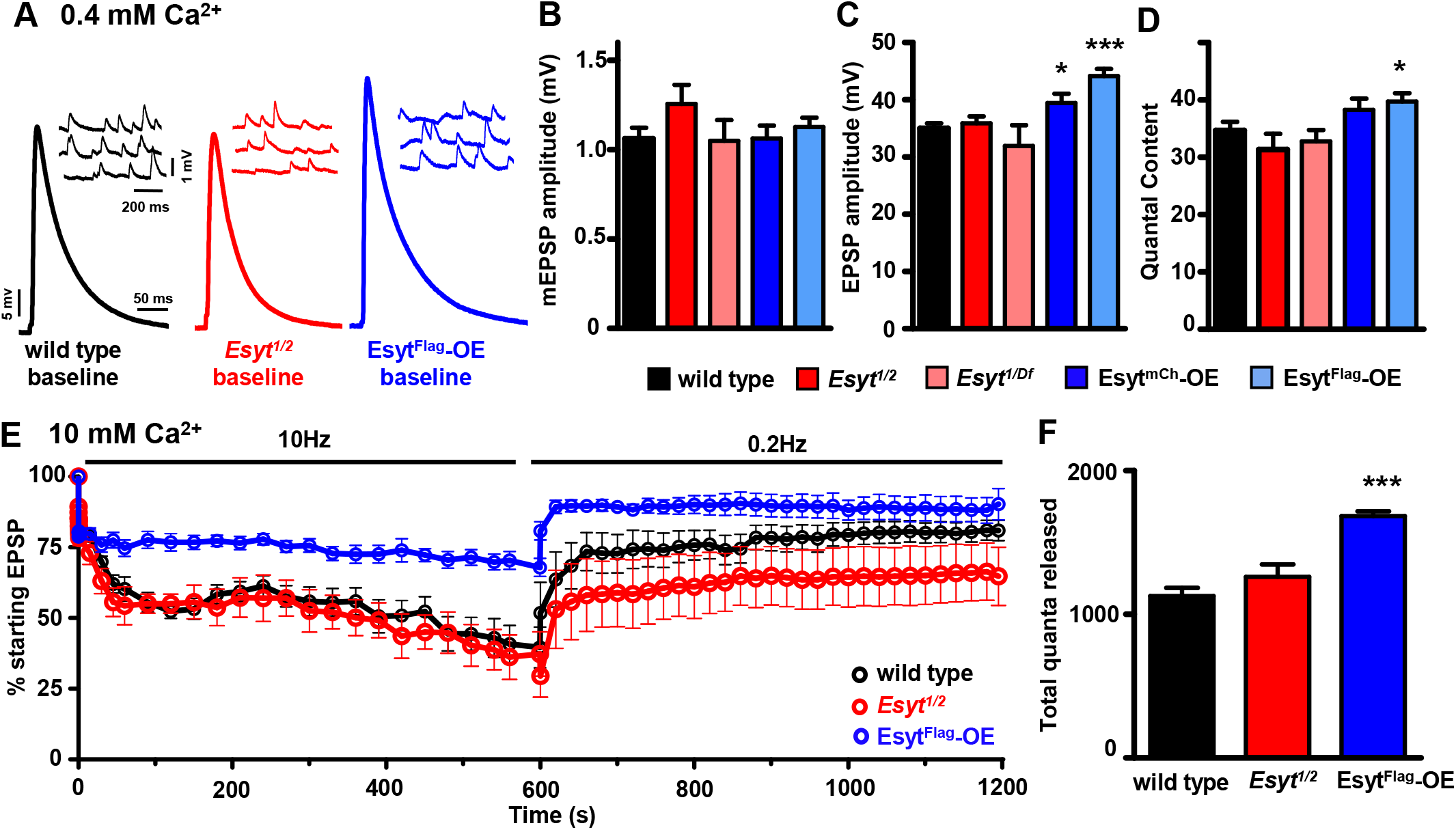
Synaptic transmission and vesicle depletion in *Esyt* mutants and Esyt-OE. (**A**) Representative electrophysiological EPSP and mEPSP traces for wild type, *Esyt* mutants (*Esyt^12^* and *Esyt^1/Df^: w;Esyt^1^/Esyt^Df^*) and Esyt-OE recorded in 0.4 mM extracellular Ca^2+^. Quantification of mEPSP amplitude (**B**), EPSP amplitude (**C**), and quantal content (**D**) of the indicated genotypes. *Esyt* mutants exhibit no significant differences in synaptic transmission compared to wild type, while Esyt-OE results in a moderate but significant increase in EPSP amplitude. (**E**) Depletion and recovery of the functional vesicle pool in the indicated genotypes. NMJs were stimulated at 10 Hz for 10 min in 10 mM extracellular Ca^2+^, then allowed to recover for 10 mins while monitoring this recovery with stimulation at 0.2 Hz. EPSP amplitudes were averaged, normalized to pre-stimulus amplitudes, and plotted as a function of time. (**F**) Quantification of total quanta released during the 10 mins of 10 Hz stimulation for the indicated genotypes. Error bars indicate ±SEM. One-way ANOVA test was performed, followed by a Tukey’s multiple-comparison test. *p≤0.05; ***p 0.001. Detailed statistical information for represented data (mean values, SEM, n, p) is shown in Table 1.

Finally, we tested whether *Esyt* was necessary to sustain synaptic transmission during elevated periods of activity. During high frequency stimulation at the *Drosophila* NMJ, endocytosis rates must be increased to sustain the elevated level of exocytosis, and any imbalance in this coupling will lead to depletion of the functional vesicle pool (Verstreken et al., 2002; Verstreken et al., 2003; Dickman et al., 2005; Haucke et al., 2011). We utilized a previously established protocol, in which we subject the NMJ to 10 Hz stimulation at elevated extracellular Ca^2+^ for 10 min, followed by recovery for an additional 10 min, taking a test pulse at 0.2 Hz (Verstreken et al., 2002; Dickman et al., 2005). This analysis revealed that wild type and Esyt-mutant synapses exhibited a similar rate of vesicle pool depletion and recovery, finishing at ~30% of the original EPSP amplitude, followed by a recovery to ~60% of the initial value (Fig 5E). In both genotypes, a similar number of total quanta were released (Fig 5F). In contrast, Esyt-OE conferred a resistance to depletion of the functional synaptic vesicle pool as well as enhanced recovery. 10 Hz stimulation of Esyt-OE NMJs revealed a slower rate of rundown and faster recovery of the depleted vesicle pool (Fig 5E), finishing at ~60% and recovering to ~90% of starting EPSP amplitudes. Indeed, more total quanta were released in Esyt-OE compared to both wild type and *Esyt* mutants during this sustained period of activity (Fig 5F). Together, this demonstrates that while the loss of *Esyt* does not impact synaptic growth, structure, transmission, or recycling, overexpression of *Esyt* in neurons promotes synaptic growth which, in turn, appears to contribute to a mild enhancement in synaptic strength while sustaining the vesicle pool during prolonged activity.

### Synaptic vesicle density and endocytosis is unchanged in *Esyt* mutants and Esyt-OE

The slowed rate of synaptic vesicle pool depletion in Esyt-OE described above could, in principle, be due to an increase in the number of synaptic vesicles participating in exocytosis and recycling at individual boutons. Alternatively, synaptic vesicle recycling at each bouton may be the same, and the increased maintenance of the vesicle pool may be due to the increased number of boutons in Esyt-OE. We therefore examined NMJ ultrastructure to determine whether a change in the density of synaptic vesicles in each bouton was apparent that may suggest an increased starting vesicle pool in Esyt-OE. We did not observe any significant change in the density or distribution of synaptic vesicles within NMJ boutons or near active zones in Esyt-OE compared with wild type and *Esyt* mutants (Fig 6A-6C). More generally, active zone length, T-bar morphology, and membrane compartments appear similar in all three genotypes (Fig 6A-6E). Thus, there is no evidence that Esyt-OE results in increased numbers or altered distribution of synaptic vesicles within NMJ boutons.

**Figure 6:**
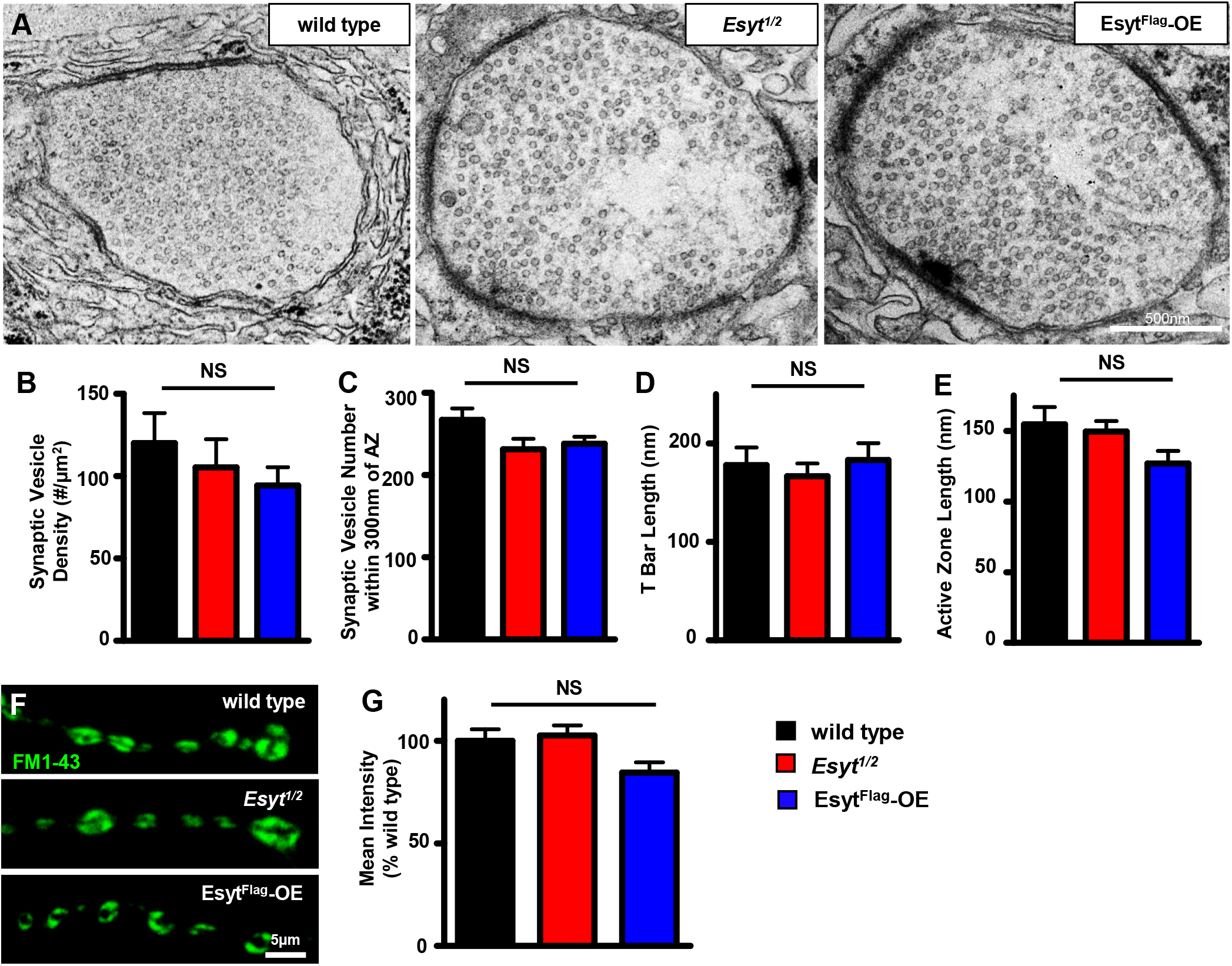
Synaptic vesicle density and endocytic pools are unchanged in *Esyt* mutants and Esyt-OE. (**A**) Representative electron micrograph images of NMJs for wild type, *Esyt* mutants, and Esyt^Flag^-OE. Quantification of synaptic vesicle density (**B**), synaptic vesicles within 300 nm of the active zone (**C**), T bar length (**D**), and active zone length (**E**) in the indicated genotypes. No significant differences were observed between genotypes. (**F**) Representative images of FM1-43 dye loading of the indicated genotypes. (**G**) Quantification of mean intensity of the FM1-43 signal in the indicated genotypes. Error bars indicate ±SEM. One-way ANOVA test was performed, followed by a Tukey’s multiple-comparison test. NS=not significant, p>0.05. Detailed statistical information for represented data (mean values, SEM, n, p) is shown in Table 1.

Despite there being no change in the number of synaptic vesicles per bouton in Esyt-OE NMJs, it is possible that more synaptic vesicles participate in exo-endocytosis during activity which, in turn, could account for the increased maintenance of the functional vesicle pool. We therefore measured the pool of vesicles participating in endocytosis during high activity using the lipophilic dye FM1-43, which is absorbed by newly endocytosed synaptic vesicles from the plasma membrane following exocytosis and is a measure of the number of vesicles participating in release at each bouton (Dickman et al., 2005; Verstreken et al., 2008; Chen et al., 2014). Following stimulation, we observed no change in the intensity or localization of the endocytic vesicle pool labeled by FM1-43 in Esyt-OE compared to wild type and *Esyt* mutants (Fig 6F and 6G). Thus we find no evidence that the number or function of synaptic vesicles per bouton is changed at NMJs in Esyt-OE. This therefore suggests that the increase in the total number of synaptic boutons, coupled with less release per bouton, together accounts for the resistance to depletion of the vesicle pool in Esyt-OE during elevated activity. Consistent with this idea, we observed a trend of reduced FM1-43 intensity per bouton following activity in Esyt-OE (Fig. 6F,G).

### *Esyt* is necessary for proper presynaptic function and short term plasticity at elevated Ca^2+^

Esyt is a putative Ca^2+^ sensor localized to axonal ER, but we were unable to observe any significant roles for Esyt in neurotransmission at lowered extracellular Ca^2+^. Indeed, in this condition, no differences in the apparent Ca^2+^ sensitivity of neurotransmission between wild type, *Esyt* mutants, and Esyt-OE were observed when quantal content was assessed across a range of lowered extracellular Ca^2+^ concentrations (Fig 7A). Interestingly, in vertebrates, Esyt1 dependent ER-PM tethering is activated only by high cytosolic Ca^2+^ concentrations (Idevall-Hagren et al., 2015; Yu et al., 2016), and we therefore hypothesized that *Esyt* function at synapses may only be revealed at elevated extracellular Ca^2+^. We therefore assayed neurotransmission and short-term plasticity at elevated extracellular Ca^2+^ concentrations (3 mM) using a two-electrode voltage clamp configuration. We observed no change in EPSC amplitude in Esyt-OE compared with wild type (Fig 7B and 7C). However, EPSC amplitudes were significantly reduced in *Esyt* mutants compared with wild type and Esyt-OE synapses, which was rescued by presynaptic expression of *Esyt* (Fig 7B and 7C). Finally, we probed short term synaptic plasticity in *Esyt* mutants by evoking four stimuli at 60 Hz in elevated extracellular Ca^2+^. Given that Esyt is a Ca^2+^ sensor localized to axonal ER, this protocol would test a role for Esyt during rapid changes in presynaptic Ca^2+^ levels (Muller et al., 2011; Muller et al., 2012; Muller et al., 2015). Wild type and Esyt-OE NMJs exhibited synaptic depression, with the fourth EPSC finishing at ~60% of the starting EPSC amplitude (Fig 7D). In contrast, *Esyt* mutants showed reduced depression, finishing at ~90% of the starting EPSC amplitude, which was rescued by presynaptic *Esyt* expression (Fig 7B and 7D). Together, this data is consistent with a Ca^2+^ sensing function of Esyt at axonal ER in promoting synaptic vesicle release during evoked activity.

**Figure 7:**
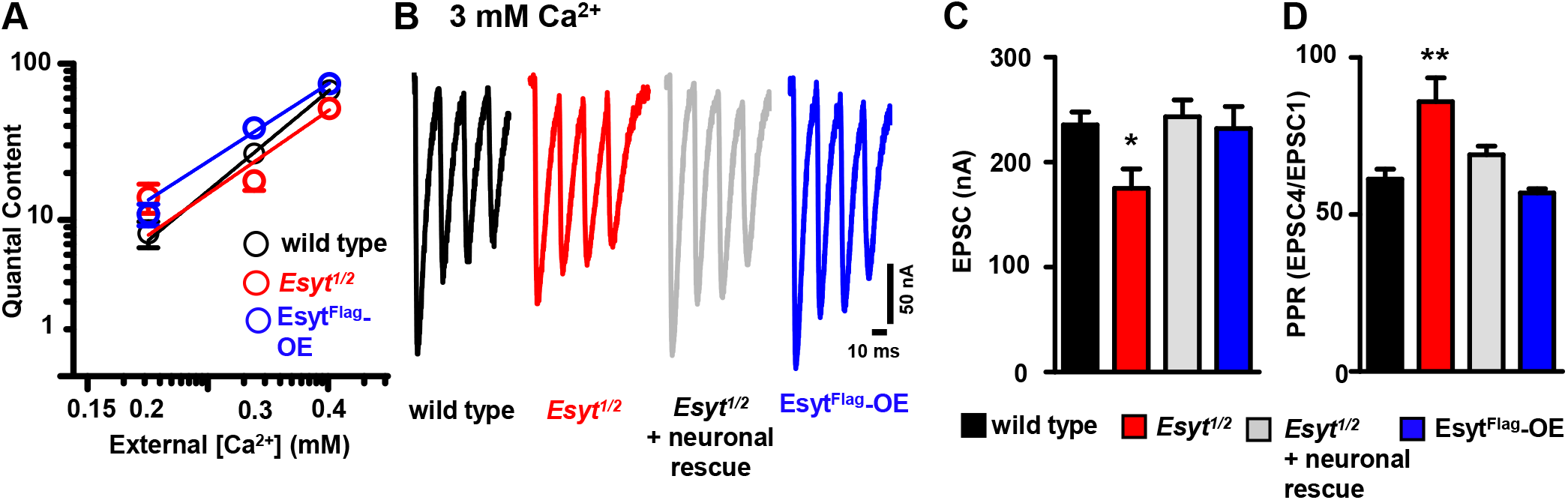
*Esyt* is required for basal neurotransmission and short-term synaptic plasticity at elevated extracellular Ca^2+^. (**A**) Quantal content plotted as a function of extracellular Ca^2+^ concentration on logarithmic scales for wild type, *Esyt* mutants (*Esyt^1/2^*), and Esyt^Flag^-OE. No significant difference was observed in the slope of the best fit lines. (**B**) Representative EPSC traces following four pulses of 60 Hz stimulation in wild type (black), *Esyt* mutants (*Esyt^1/2^*, red), *Esyt* neuronal rescue (*w;OK6-Gal4/UAS-Esyt^Flag^; Esyt^1^/Esyt^2^*), and Esyt^Flag^-OE (blue) in 3 mM extracellular Ca^2+^. (**C**) Quantification of average EPSC amplitude in the indicated genotypes. (**D**) Quantification of average EPSC ratio (4^th^ EPSC/1^st^ EPSC) for the indicated genotypes. *Esyt* mutants show reduced neurotransmission and short-term synaptic plasticity at high extracellular Ca^2+^. Error bars indicate ±SEM. One-way ANOVA test was performed, followed by a Tukey’s multiple-comparison test. *p≤0.05; **p 0.01. Detailed statistical information for represented data (mean values, SEM, n, p) is shown in Table 1.

### *Esyt* has no role in presynaptic homeostatic potentiation

Thus far, we have found that *Esyt* has no apparent role in synaptic growth and structure, but is required to promote synaptic vesicle release at elevated extracellular Ca^2+^. Interestingly, elevated Ca^2+^ influx at presynaptic terminals at the *Drosophila* NMJ has been demonstrated to drive an adaptive form of synaptic plasticity referred to presynaptic homeostatic potentiation (PHP) (Muller and Davis, 2012; Davis and Muller, 2015). At this synapse, pharmacological or genetic perturbations to postsynaptic glutamate receptors triggers a precise retrograde increase in presynaptic release that compensates for reduced receptor functionality, restoring baseline levels of synaptic strength (Frank, 2014). We hypothesized that Esyt may be an axonal ER Ca^2+^ sensor that promotes vesicle release in response to elevated presynaptic Ca^2+^ influx during PHP expression. We assayed acute PHP in *Esyt* mutants and Esyt-OE. Application of the glutamate receptor antagonist philanthotoxin-433 (PhTx; (Frank et al., 2006)) to wild-type NMJs led to the expected ~50% reduction in mEPSP amplitude but normal EPSP amplitude because of a homeostatic increase in quantal content (presynaptic release) (Fig 8A and 8B). Similarly, PhTx reduced mEPSP amplitudes in both *Esyt* mutants and Esyt-OE, and both genotypes exhibited a robust increase in quantal content (Fig 8A and 8B). Thus, loss or increased expression of *Esyt* has no consequence on the acute induction or expression of PHP.

**Figure 8:**
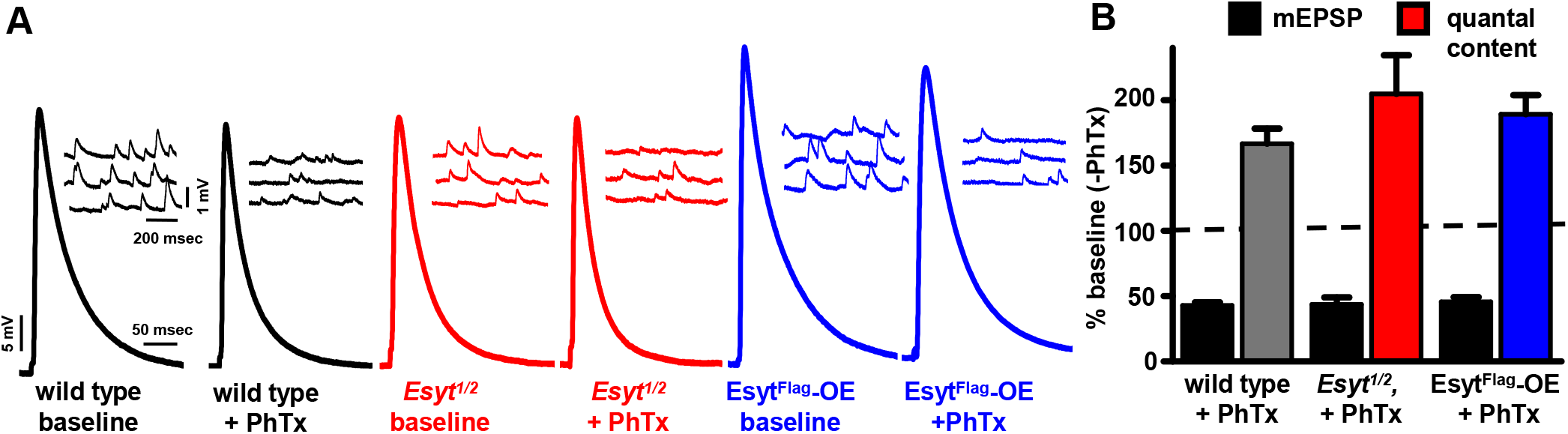
*Esyt* is dispensable for presynaptic homeostatic plasticity. (**A**) Representative electrophysiological EPSP and mEPSP traces for wild type, *Esyt* mutants (*Esyt^1/2^*), and Esyt^Flag^-OE before (baseline) and following PhTx application. Note that while mEPSP amplitudes are reduced following PhTx application, EPSP amplitudes recover to baseline levels because of a homeostatic increase in presynaptic release (quantal content). (**B**) Quantification of mEPSP and quantal content values after PhTx application normalized to baseline values. No significant differences were observed. Error bars indicate ±SEM. One-way ANOVA test was performed, followed by a Tukey’s multiple-comparison test. Detailed statistical information for represented data (mean values, SEM, n, p) is shown in Table 1.

## DISCUSSION

We have generated the first mutations in the single *Drosophila Esyt* ortholog and characterized the presynaptic localization and functions of this gene at the NMJ. We demonstrate that *Drosophila* Esyt is localized to an extensive axonal ER network. Although Esyt was previously shown to mediate ER-PM tethering and to promote lipid exchange between the two membranes in non-neuronal cells, we find no evidence that lipid balance is altered at *Esyt* mutant synapses. Further, synaptic transmission and homeostatic plasticity are surprisingly unperturbed in the absence of *Esyt* at lowered extracellular Ca^2+^ and during sustained levels of synaptic activity. In contrast, presynaptic overexpression of *Esyt* promotes synaptic growth and, in turn, synaptic transmission during elevated activity. Finally, we reveal an important function for *Esyt* in facilitating presynaptic release at elevated Ca^2+^ levels. Together, our study establishes Esyt as an ER-localized C2 domain protein that regulates synaptic growth when overexpressed and neurotransmission at elevated Ca^2+^ levels.

### *Drosophila* Esyt is dispensable for many synaptic functions

*Esyt* is evolutionarily conserved from yeast to humans (Min et al., 2007; Manford et al., 2012), suggesting this gene family subserves important and fundamental functions that have been selected for and maintained throughout evolution. Studies in yeast and mammalian cell culture revealed that Esyt mediates lipid transfer at ER-PM contact sites in a Ca^2+^-dependent manner (Saheki et al., 2016; Yu et al., 2016; Saheki and De Camilli, 2017), raising the intriguing possibility that Esyt may modulate lipid metabolism during growth and activity at synapses. However, recent studies in which all three *Esyt* genes are genetically mutated in mice have failed to find any apparent defects in viability, fertility, synaptic or ER protein composition, nor in ER/mitochondrial stress responses (Sclip et al., 2016; Tremblay and Moss, 2016). Consistent with these studies, we find that *Esyt* is surprisingly dispensable for lipid balance and homeostasis at presynaptic terminals, and that synaptic growth, function, and plasticity appear relatively unperturbed at the *Drosophila* NMJ. We consider two possibilities, not mutually exclusive, to explain why synapses robustly develop and function in the absence of *Esyt*.

First, Esyt may participate in such fundamental and essential processes that organisms may in turn have evolved multiple redundant mechanisms to ensure robustness in these critical pathways. Indeed, in pioneering work in yeast, the *Esyt* orthologs tricalbins mediate ER-PM tethering, but this tethering is only disrupted upon loss of three additional ER-PM proteins, Ist2, Scs2, and Scs22, in addition to the three tricalbins (Manford et al., 2012). This suggests some level of functional redundancy within and beyond Tricalbin isoforms. Further, a recent study demonstrated an apparent compensatory increase in expression of the ER-PM tethering proteins Orp5, Orai1, and TMEM110 in Esyt1, 2, and 3 mutant mice (Tremblay and Moss, 2016). In *Drosophila*, a number of proteins are localized to ER and have been proposed to contribute to lipid balance and homeostasis in other cellular compartments (Wenk et al., 2003; Carrasco and Meyer, 2011; Hammond et al., 2014; Dickson et al., 2016). Thus, there is substantial evidence for a variety of proteins at the ER and other compartments having potentially redundant function with Esyts.

Second, Esyt may only have functions revealed in a specific context that was not tested in our study. We find that Esyt is not essential to maintain lipid homeostasis at synapses, at least for the major phospholipids PI(4,5)P2 and PI(3)P. This demonstrates that lipid balance and membrane homeostasis can be maintained during the extreme demands of regulated membrane trafficking and exchange at presynaptic terminals in the absence of *Esyt*. In retrospect, this may not be surprising, as a lipid cycle nested within the synaptic vesicle cycle has long been known to exist at synapses, supported by key synaptic proteins such as Synaptojanin, Rab5, Minibrain kinase/Dyrk1A, and Sac1 (De Camilli et al., 1996; Nemoto et al., 2000; Wenk and De Camilli, 2004; Chen et al., 2014). Importantly, there is no known involvement or requirement for acute lipid transfer from the ER in synaptic vesicle recycling (De Camilli et al., 1996; Wenk and De Camilli, 2004). Accordingly, lipid homeostasis during synaptic vesicle trafficking, like protein homeostasis, may be sufficiently embedded and coupled in membrane trafficking itself so as not to lead to imbalances, even during rapid exo- and endocytosis. Thus, a variety of redundant, tissue-specific, and/or specialized functions of *Esyt* likely explains the relatively subtle effects due to loss of this fundamental gene.

### Esyt localizes to axonal ER and promotes neurotransmission at elevated Ca^2+^ levels

We find that Esyt is localized to the axonal ER. Further, we demonstrate that Esyt promotes neurotransmission, but only at elevated Ca^2+^ levels. These findings suggest that Esyt may promote presynaptic function by coupling local Ca^2+^ dynamics to mechanisms that modulate intracellular Ca^2+^ release from the axonal ER. Indeed, axonal ER has emerged as a crucial organelle that can sense and respond to dynamic change in cytosolic Ca^2+^ to modulate presynaptic function and plasticity (Verkhratsky, 2005; Bardo et al., 2006; Kwon et al., 2016; de Juan-Sanz et al., 2017). For example, Ca^2+^ influx from the extracellular space can induce additional Ca^2+^ release from the ER via ryanodine receptors, a process referred as Ca^2+^-induced Ca^2+^-release (CICR), which can be activated during single action potentials or during the trains of stimuli (Verkhratsky, 2005; Bardo et al., 2006; Kwon et al., 2016; de Juan-Sanz et al., 2017). While the molecular mechanism for how Esyt promotes neurotransmission is unclear, it is unlikely to be through developmental alterations in synaptic structure, since we did not find any defects in these processes in *Esyt* mutants. Rather, Esyt likely has a role at axonal ER in acutely modulating presynaptic function during activity. An intriguing possibility is that Esyt may work in conjunction with ryanodine receptors as Ca^2+^ sensors that promotes CICR at the presynaptic terminal in response to activity. In this model, Esyt may respond to elevated Ca^2+^ at synapses during single action potentials, leading to a supplemental source of presynaptic Ca^2+^, perhaps through release of intracellular ER stores.

### Esyt, axonal ER, and synaptic growth

Perhaps the most striking and unexpected finding was that elevated expression of *Esyt* in motor neurons led to increased synaptic growth and a moderate increase in neurotransmission. As discussed above, we find no evidence that Esyt is involved in the regulated membrane trafficking pathway driving synaptic vesicle recycling. It is therefore attractive to consider that elevated *Esyt* expression at axonal ER promotes increased membrane trafficking to the plasma membrane through the constitutive pathway. Indeed, the membrane necessary for synaptic growth is delivered through the constitutive membrane trafficking pathway, as synapses are able to form and grow even when toxins are expressed that block or inhibit synaptic vesicle fusion (Broadie et al., 1995; Sweeney et al., 1995; Dickman et al., 2006; Choi et al., 2014). Of course, the ER is well established to be a key node in constitutive membrane trafficking to the PM (Aridor and Fish, 2009; Pfenninger, 2009; Ramirez and Couve, 2011), and increased *Esyt* expression at axonal ER may facilitate the rate of membrane delivery to the presynaptic terminal. This, in turn, may lead to excess synaptic growth, as observed in other mutants with excess presynaptic membrane (Koh et al., 2004; Dickman et al., 2006; O’Connor-Giles et al., 2008). Thus, the unanticipated finding that increased *Esyt* expression promotes synaptic growth raises intriguing possibilities for axonal ER perhaps being a rate-limiting step in membrane delivery at synapses.

Our results establish Esyt as an axonal ER Ca^2+^ sensor that promotes presynaptic function at elevated extracellular Ca^2+^ levels and may also have unanticipated roles in synaptic growth. Further work is needed to elucidate the molecular mechanism of how Esyt couples cytosolic Ca^2+^ to presynaptic ER functions to ultimately modulate transmission and signaling. While axonal ER was first observed over 40 years ago (Tsukita and Ishikawa, 1976; Ramirez and Couve, 2011), the functions of this complex organelle has remained enigmatic. Recent studies have begun to reveal how axonal ER sculpts presynaptic Ca^2+^ dynamics and modulates presynaptic function and plasticity (de Juan-Sanz et al., 2017), roles that seem certain to contribute to a variety of neurological diseases.

## ACKNOWLEDGEMENTS

We acknowledge the Bloomington Drosophila Stock Center and the Iowa Developmental Studies Hybridoma Bank for genetic and antibody reagents. We also thank Christopher Buser at Droseran LLC (Pasadena, CA) for technical assistance with electron microscopy. This work was supported in part by a USC Provost Fellowship to KK and by an NIH grant to DKD (NS019546), as well as funding from the Alfred P. Sloan, Ellison Medical, Mallinckrodt, Klingenstein-Simons, and Whitehall Foundations to DKD.

## AUTHOR CONTRIBUTIONS

KK and DKD designed the research and KK, DK, DS, and XL performed experiments. KK, DK, DS, and XL analyzed data. KK and DKD wrote the manuscript.

The authors declare no competing financial interests.

## FIGURE LEGENDS

**Table 1: Absolute and additional values for normalized and presented data**. The figure and panel, genotype, and conditions used are noted (external Ca^2+^ concentration as well as whether PhTx was applied or not). Average values with standard error values noted in parentheses are shown for all morphology data such as neuronal membrane surface area, bouton number, as well as intensity values for all represented synaptic proteins and postsynaptic glutamate receptors are shown. For electrophysiological experiments, all passive membrane properties (input resistance, resting membrane potential), and mEPSP, EPSP, QC, number of data samples (n) and p-values from the statistical test are shown.

